# Principal component analysis revisited: fast multi-trait genetic evaluations with smooth convergence

**DOI:** 10.1101/2024.06.06.597390

**Authors:** Jon Ahlinder, David Hall, Mari Suontama, Mikko J Sillanpää

## Abstract

A cornerstone in breeding and population genetics is the genetic evaluation procedure, needed to make important decisions on population management. Multivariate mixed model analysis, in which many traits is considered jointly, utilizes genetic and environmental correlations between traits to improve the accuracy. However, the number of parameters in the multi-trait model grows exponentially with the number of traits which reduces its scalability. Here, we suggest using principal component analysis (PCA) to reduce the dimensions of the response variables, and then using the computed principal components (PC) as separate responses in the genetic evaluation analysis. As PCs are orthogonal to each other, multivariate analysis is no longer needed and separate univariate analyses can be performed instead. We compared the approach to traditional multivariate analysis in terms of computational requirement and rank lists according to predicted genetic merit on two forest tree datasets with 22 and 27 measured traits respectively. Obtained rank lists of the top 50 individuals were in good agreement.

Interestingly, the required computational time of the approach only took a few seconds without convergence issues, unlike the traditional approach which required considerably more time to run (seven and ten hours respectively). Our approach can easily handle missing data and can be used with all available linear mixed models software as it does not require any specific implementation. The approach can help to mitigate difficulties with multi-trait genetic analysis in both breeding and wild populations.

## Introduction

Phenotyping is a critical process in any breeding program with the aim to improve the genetic level of the traits of interest. By accurately characterizing the traits, breeders can make informed decisions about which individuals to select in breeding populations to achieve expected increase in genetic merit shown as increased productivity, quality, vitality depending on the breeding objective. In the near future, many novel high-throughput phenotyping techniques could transform how traits are defined and recorded and could easily reach thousands. For example, remote sensing techniques, such as LiDAR (Light Detection and Ranging), which can capture detailed 3D structural information about crops and trees including canopy height, width and architecture, disease status and condition to name a few [42]. Another high-throughput example involves gene expression data, where linear mixed effect models (LMM) have been used to identify sources of variation in human medicine studies of HIV infection [85], identifying genotype-by-environment (GxE) interactions in body mass index [57] and human brain regions [77]. In a plant breeding application, [72] showed how to use LMM to jointly analyze grain yield and hyperspectral reflectance traits measured in wheat (*Triticum aestivum*) field trials.

Multi-trait LMM analysis was introduced in quantitative genetics by [36] and encompass both genetic covariance component estimation and estimation of breeding values (EBV). Compared to analyzing each trait separately, the advantages of multi-trait analysis are:

- increased prediction accuracy of breeding values for un-phenotyped individuals,
- increased statistical power as available data is more efficiently used,
- increased parameter estimation accuracy by exploiting correlations between traits.

In particular, multi-trait LMM analysis can provide more accurate estimations in the case of traits with a low heritability (i.e. a proportion of trait variation attributable to genetic factors), populations of small size or if missing data is present [32, 65]. Accurate estimation of variance components and functional parameters, such as heritabilities and genetic correlations, is important because prediction error variances for estimated random effects increase as the differences between estimated and true values of variance components increase [59]. Many studies have been published comparing the performance of single and multiple-trait LMMs. For example, [3] compared EBVs obtained from both multi-trait and single-trait LMMs via the best linear unbiased predictor (BLUP) for tree height, DBH, and tree volume in *Eucalyptus ssp*. and predicted higher selection response with the multi-trait BLUP analysis. Using simulations, [32] showed that for traits with missing data, the EBVs obtained in the multiple-trait analysis resulted in more reliable genomic predictions.

Unfortunately, the number of parameters in multi-trait LMMs grows exponentially with the number of traits due to added pair-wise correlation parameters, and the required computational effort therefore grows even more because of the need to invert a (likely) large coefficient matrix at each iteration in the inference procedure, at least for most available algorithms, such as restricted maximum likelihood (REML) [63] or Bayesian blocked Gibbs sampling [78]. For example, often various convergence problems arises, and this can lead to unstable parameter estimates [43, 56]. In most practical applications, only a few traits can be analysed simultaneously, which is not optimal as shared information via correlations is not used efficiently in the inferential procedure, causing biased parameter estimates, both for location (i.e. breeding values) and scale (covariance components and heritability). Methods that can circumvent this problem would be sought after.

A number of alternative approaches that tries to circumvent the problems of standard multi-trait LMM analysis have been suggested in the literature. [47] suggested the use of reducing the rank of the covariance matrix by principal component analysis (PCA) or by factor analytic (FA) models to improve multi-trait LMM analysis. By directly estimate the leading principal components, most of the important information is kept while reducing the computational burden to estimate the covariance matrices [47, 53, 54]. This can make the model easier to estimate and interpret, especially in advanced breeding programs where the full covariance structure is complex. [53] showed how this approach could be used for selection and multi-trait LMM analysis of carcass traits for Angus cattle, where she suggested that the first seven of the PCs were sufficient to obtain estimates of breeding values without loss in the expected accuracy of evaluation. The approach has recently been shown to reduce computational burden in dense genomic marker-derived covariance matrices by [55]. However, a drawback is that there is an obvious loss of information when the rank of the covariance matrix is reduced, information that could be be important for some of the traits included.

The use of dimension reduction techniques to simplify the covariance structure have been a popular choice in crop and forest tree breeding when estimating GxE interactions in multi-trait LMM evaluations [9, 12, 50, 66, 73]. In a review of GxE in forest tree breeding, [50] reviewed analytical methods for inferring G×E effects, including factor analytic modelling, and its application in analysis of field trials of forest tree species including Pine spp, Eucalypt spp, Spruce spp, and Poplar spp. [12] incorporated factor analysis to reduce the rank of the covariance matrix which enabled the incorporation of 19 traits simultaneously into the multi-environment LMM analysis of Scots pine (*Pinus sylvestris*) field trials. As a result, they found that the main driver of detected GxE was differences in temperature sum among trial sites. [67] analyzed the mean annual height increment, the mean annual diameter increment, and wood density in a series of field trials of European larch using FA models: the inferred genetic correlations between sites showed low to high GxE, with growth traits exhibited more GxE than wood density.

Another popular approach in various breeding scenarios is to perform canonical transformation to improve the performance of multi-trait LMMs [20, 40, 84], in which a matrix decomposition technique is applied on both genetic and residual covariance matrices. After the transformation is applied, BLUP values can be computed for each trait using univariate LMMs. Then the obtained solution can be back transformed to the original scale, which thereby facilitates interpretation. Unfortunately, a typical requirement for canonical transformation is that covariance matrices either need to be known or ad-hoc estimated before the transformation: this limits the usefulness of the approach as uncertainties in the estimation procedure is not accounted for.

Instead of simplifying the covariance structure, a more direct approach would be to consider transforming the phenotypic traits. The use of PCA to simplify multi-trait LMM analysis by operating on the phenotypic trait records is not new and have previously been used to perform genetic variance component and heritability estimation [5, 38]. [14] used PCA of skeletal variation in a population of Portuguese water dogs to reveal groups of traits defining skeletal structures and associate it with quantitative trait loci (QTLs). A related PCA based approach has been proposed for linkage analysis [62] and genome-wide association analysis [4, 86]. The advantages in breeding value and genetic variance component inference of using PCA is that it can handle a large set of traits by transforming them into orthogonal principal components (PC) which can be seen as trait combinations with similar characteristics that cannot be measured directly. As each PC is orthogonal, they can be analyzed independently with univariate models. This procedure would be very fast and converge very quickly as opposed to multivariate analysis of a large set of traits, especially when dealing with unbalanced longitudinal data [1] or with large sets of predictors [51]. One problem with this approach is that it cannot handle missing data, at least not the standard single value decomposition (SVD) approach, which restrict its use in general breeding applications. Thus, more effort into using PCA directly on the phenotypic profiles which includes missing data is warranted.

A similar approach using factor analytic modelling, which operates on the response matrix, have recently been proposed [71, 72]. By introducing latent variables via a mixed effect factor model, all sources of correlation among the traits can be accounted for and corresponding univariate independent LMMs could be analyzed. With this approach, MegaLMM [72], three plant breeding data sets with tens-of-thousands of traits were analyzed, and obtained results showed improved prediction accuracy of genetic values and improved computational speed compared to results obtained by traditional methods. As a model-based approach it can handle missing data, but it needs a special software implementation which limits its general use.

Here, we aim at improving (co)variance component and breeding value estimation of large scale phenotyping efforts by re-introducing PCA dimension reduction technique to obtain reduced space traits. This suggested method can easily be analysed with standard univariate LMMs. In doing so, we circumvent the problems of convergence in the REML analysis to estimate covariance components, at a fraction of the required computational time, compared to a multivariate analysis. A novelty in this proposed approach is how missing data can be handled efficiently in the ordination step by utilizing a model based PCA for imputation. Two typical forest tree data sets are used to highlight the performance of the approach: one Scots pine field trial included in the north Swedish breeding program and one Loblolly pine breeding population with traits scored in several trials. Because all data was pre-adjusted for trial specific design and environmental effects, the continuous nature of the adjusted data facilitated the PCA analysis.

## Materials and methods

### Multi-trait linear mixed effect model

Under Gaussian assumptions, the multivariate version of the linear mixed-effect model (LMM) can be written as:

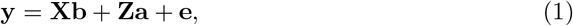

where **y** is the response vector containing each of *m* traits represented sequentially for *n* individuals in a single vector. This is obtained by taking vec-operation of the multivariate observation matrix of dimension *n* × *m*. **X** is a *p* × *nm* design matrix for fixed effects (with ones, zeros or regression measurements as their elements) in *p* fixed effects in *m* traits. This is a block-matrix with *m* blocks of size *p* × *n*. Similarly, **b** represents the fixed-effects coefficient vector with dimension *pm*, **Z** is the design matrix for random effects with dimension *n* × *m*, **a** denotes the random-effects vector (i.e. polygenic additive effects) with dimension *nm*, **e** represents the error vector (i.e. residuals) of size *nm*. Furthermore, the random effects are assumed to follow a multivariate normal distribution **a** ∼*MV N*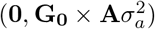, where **G**_**0**_ is a *m* × *m* genetic correlation matrix, **A** is a *n* × *n* additive genetic relationship matrix, and 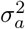 is the additive genetic variance component. The residuals are assumed to be multivariate normally distributed as **e** ∼*MV N*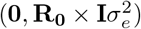, where *R*_0_ is a *m* × *m within individual* residual covariance matrix and *I* is an *n* × *n* identity matrix, viz., *between individual* residual covariance matrix and 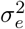 is the residual variance component.

### Standard singular value decomposition based PCA

PCA reduces the dimensionality of data while preserving its essential information [37, 44, 82]. PCA is computed for *n* × *m* multivariate observation matrix **Y**, where *n* is the number of individuals and *m* traits. If *n* ≤ *m*, it is practical to calculate it for *n* × *n* matrix of **YY**^*′*^. Otherwise, it is calculated for *m* × *m* matrix of **Y**^*′*^**Y**. Let us represent a scaled symmetric covariate matrix **YY**^*′*^ as a product of two orthogonal lower triangular matrices **Q** and one diagonal matrix **D** such as **YY**^*′*^ = **QDQ**^*′*^.

Orthogonality means that **Q**^*′*^**Q** = **I**. Now, if we multiply both sides from left and right with **Q**^*′*^ and **Q**, respectively, we obtain **Q**^*′*^**YY**^*′*^**Q** = **D** = *diag*(*λ*_1_, *λ*_2_, .., *λ*_*p*_), where the right hand side of equation contains eigenvalues of matrix **YY**^*′*^ in the diagonal in the ascending order *λ*_1_ ≥*λ*_2_ ≥..≥ *λ*_*p*_. This is closely related to so called singular value decomposition (SVD) and also to Cholesky decomposition [31]. For PCA, the singular values are the square roots of the eigenvalues of the covariance matrix, and both eigenvalues and singular values provide insights into the phenotypic variability and importance of different components (eigenvectors or basis vectors) in transforming and summarizing the observed phenotypic profiles.

### Model-based PCA

PCA is a linear transformation of the covariance matrix of the data to the space where different directions are independent. As an alternative to the algorithmic-based exact PCA is to fit the transformation model to the data statistically using a probabilistic model based approach (i.e. observed data points are generated from a probabilistic distribution) [76]. This of course requires distributional model assumptions which increase transparency but makes the transformation somewhat noisy. One advantage is the possibility to include handling of missing values as part of the hierarchical model. [60] suggested a Bayesian model based version of PCA, or BPCA, which simultaneously fits a probabilistic model and infer latent variables (i.e., principal components, PC). The main step that include the missing value imputation in BPCA is the PC regression step: for the i:th trait 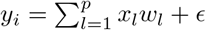 where *w*_*l*_ is the l:th principal axis vector and *x*_*l*_ is the linear coefficient to be estimated (also called the factor score), *p* is the total number of components used, and *ϵ* is the residual. The goal is to minimize the sum of squared errors ∥*ϵ* ∥^2^ for *Y* (i.e. for all traits) by using PCA. As we have missing data, the principal axis matrix *W* can be divided into a complete and missing data part: *W* = (*W* ^*obs*^, *W* ^*miss*^). The factor scores *x* are then obtained by minimizing the residual error for the observed data *y*^*obs*^, and then used to obtain *y*^*miss*^ = *W* ^*miss*^*x*. [60] used Bayesian inference, via a variational Bayes algorithm [6] to estimate model parameters and missing records. Interested readers are invited to see [60] for further details of the missing data imputation steps.

### Implementation details

Standard SVD-based PCA were performed using the prcomp function from the R package stats [68] with default settings. We used the R package pcaMethods [74] to apply BPCA to impute missing data and perform ordination by using the pca function with parameters maxSteps = 10000 and threshold = 1e-05. In addition, we also used the missForest R package [75] as a comparison of the effect of missing data imputation. ASReml-R [11] was used to infer genetic parameters both in the standard bivariate approach and in the univariate analysis of PCs, with the workspace parameter increased to 4096mb, and the aising parameter set to true. To infer the association between selection index and PCs, we used Lasso *L*_1_ regularization via the glmnet R package [25]. The strength of the penalty applied to the coefficients (i.e. *λ*) was set to the value 0.15 in order to keep only a few PCs per index. To compare ordinations (i.e. loadings) obtained with SVD-PCA and BPCA on imputed data, Eucledian distance of the loading matrices were first calculated (dist function) and then compared using a Mantel test (mantel function) available in the Vegan R package [61]. Correlations between EBV obtained with SVD-PCA and BPCA, and with imputed and original data was calculated via the cor.test function using Pearson’s product moment correlation in R. Rank lists were compared using association test between paired samples with Kendall’s *τ* method implemented in the cor.test function. The null-hypothesis tested was if the true tau was equal to 0 (i.e. no association). All LMM software’s tested on the first three PCs of the Loblolly pine data were used with default settings. To perform clustering analysis of the loadings, Ward’s method was used in conjunction with Euclidean distance via the hclust function in R, stats package. All figures were produced using the ggplot2 R package [81].

To calculate standard deviation of the narrow-sense heritability based on estimated variance components and their standard deviations (StdDev), we made use of the following Taylor’s approximation: StdDev 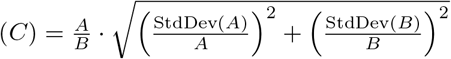 assuming absence of co-variation between *A* and *B*, where in our case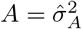, and 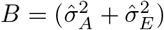. All StdDevs were estimated in respective software.

For further details including R code and data, please visit https://github.com/jonhar97/Reduced_phenotype_MME.

### Analysed Scots pine data

The Scots pine (*Pinus sylvestris L*.) field trial was designed to test the performance of available genotypes in seed orchard 412 Domsjöänget. The trial was established in 1971, located in Vindeln, Sweden 64.18 ° N, 19.34 ° E and consisted of 206 full-sib families obtained from controlled crosses of 52 seed orchard parents and five local stand seed sources, totalling 8160 plants at 3.95 hectare of land. The plants where spaced at 2.2 × 2.2 meter squares in single tree plots. The trees were measured after 10, 14, 26, and 47 growing seasons for production and quality related traits (Table 1). The trial was thinned after the 26 year measurement. Previous studies have reported moderate heritability estimates for tree height and diameter [23, 24, 34]. We used two alternative selection indices with equal weight to all included traits at age 26 (i.e. close to final evaluation of the trial in north of Sweden):

- production using height (Hjd 26) and diameter (Dia 26)
- production and tree stem quality using equal weights for height (Hjd 26), diameter (Dia 26), branch angle (Gvin 26), and (negative) branch diameter (Gdia 26)

**Table 1.** Scots pine trait statistics. Traits measured in the Scots pine progeny trial. Standard errors are within parenthesis. Estimates of narrow-sense heritabilities for the 1685 and 6044 tree subsets are denoted as 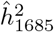 and 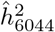, respectively.

| Trait | Age | Trait type | Obs. | $\hat{h}_{1685}^2$ | $\hat{h}_{6044}^2$ | Additional info |
| --- | --- | --- | --- | --- | --- | --- |
| Dia_14 | 14 | Production | 2765 | 0.077 (0.035) |  | Diameter at breast height |
| Dia_26 | 26 | Production | 5244 | 0.120 (0.041) | 0.147 (0.032) | Diameter at breast height |
| Dia_47 | 47 | Production | 4302 | 0.242 (0.055) | 0.208 (0.040) | Diameter at breast height |
| Ftop_47 | 47 | Quality | 4425 | 0.047 (0.028) | 0.041 (0.017) | Number of top shoots |
| Gant_14 | 14 | Quality | 2767 | 0.236 (0.055) |  | Number of branches per whorl |
| Gdia_26 | 26 | Quality | 5313 | 0.372 (0.067) | 0.320 (0.052) | Average branch diameter |
| Gdiagr130_14 | 14 | Quality | 2151 | 0.131 (0.045) |  | The diameter of the largest branch |
| Gvin_26 | 26 | Quality | 5313 | 0.382 (0.066) | 0.347 (0.052) | Branch angle |
| Gvingr130_14 | 14 | Quality | 2148 | 0.479 (0.071) |  | The angle of the largest branch |
| Hjd_10 | 10 | Production | 6027 | 0.112 (0.040) | 0.191 (0.037) | Total tree height |
| Hjd_14 | 14 | Production | 5506 | 0.262 (0.057) | 0.249 (0.043) | Total tree height |
| Hjd_26 | 26 | Production | 5248 | 0.573 (0.072) | 0.369 (0.055) | Total tree height |
| Hjd_47 | 47 | Production | 4023 | 0.390 (0.066) | 0.323 (0.052) | Total tree height |
| Sprant_14 | 14 | Quality | 2898 | 0.084 (0.035) |  | Top shoot count |
| Sprant_26 | 26 | Quality | 5316 | 0.110 (0.040) | 0.147 (0.032) | Top shoot count |

To pre-adjust phenotypic records, we followed e.g. Calleja-Rodriguez2019 by using the following set of predictors:

- fixed effect: intercept,
- random effects: plot, rows within plot, columns within plot,
- residual covariance structure: AR1 autocorrelation term on rows and columns.

### Analysed Loblolly pine data

The loblolly pine (*Pinus taeda L*.) breeding population dataset was published by [69], which originated from controlled crosses of 32 parents (22 field-selected F0 plus trees and 10 selected F1 progeny) representing a wide range of accessions from the southeastern USA. A subset of 926 genotypes of the F2 offspring was selected for extensive phenotyping in three replicated studies for growth, developmental, and disease-resistance traits measured at one, two, three, four and six years 2. We defined selection index inspired by [39]:

- production index with equal weights on EBVs for height and diameter at age 6
- production and disease susceptibility index with EBVs for height, diameter, and (negative) rust infection at equal weight
- production and wood quality with EBVs for height, diameter, stiffness, and density at equal weight

## Results

### PCA based multi-trait LMM analysis of a Scots pine field progeny trial

#### Accurate EBV ranking

Two subsets were extracted from the original data of 8,100 trees to check the robustness and performance of the PCA based method: one smaller subset of 1685 tress scored for 15 traits without missing data, and a larger subset of 6044 trees measured with 10 traits with 13.3 % of missing data (Supplementary figure S1, Table 1). Prior to the analysis, all data were spatially adjusted using an first order spatial autoregressive (AR1) model as suggested by [22] to remove micro-environmental variation. First, we focus on the analysis of the complete 15 trait subset to compare the approaches.

The PCA revealed strong clustering tendencies among the recorded traits, where the first PC differentiated between production and outer tree quality traits, while retaining a large chunk of the total phenotypic variation (Fig 1). In total, 15 PCs were used to explain all phenotypic variation, where the first three components explained 35.6, 13.7 and 8.8 % of the variation (cumulative: 58.1 %). This manifested as a strong linear correlation between the first component and multiple production traits such as height and diameter at different ages. To confirm the grouping of the traits using the obtained loadings in the PCA, we performed hierarchical clustering of the loadings which showed the same partition of traits into production and quality groups divided at the highest hierarchy level (Supplementary figure S2).

**Figure 1.**
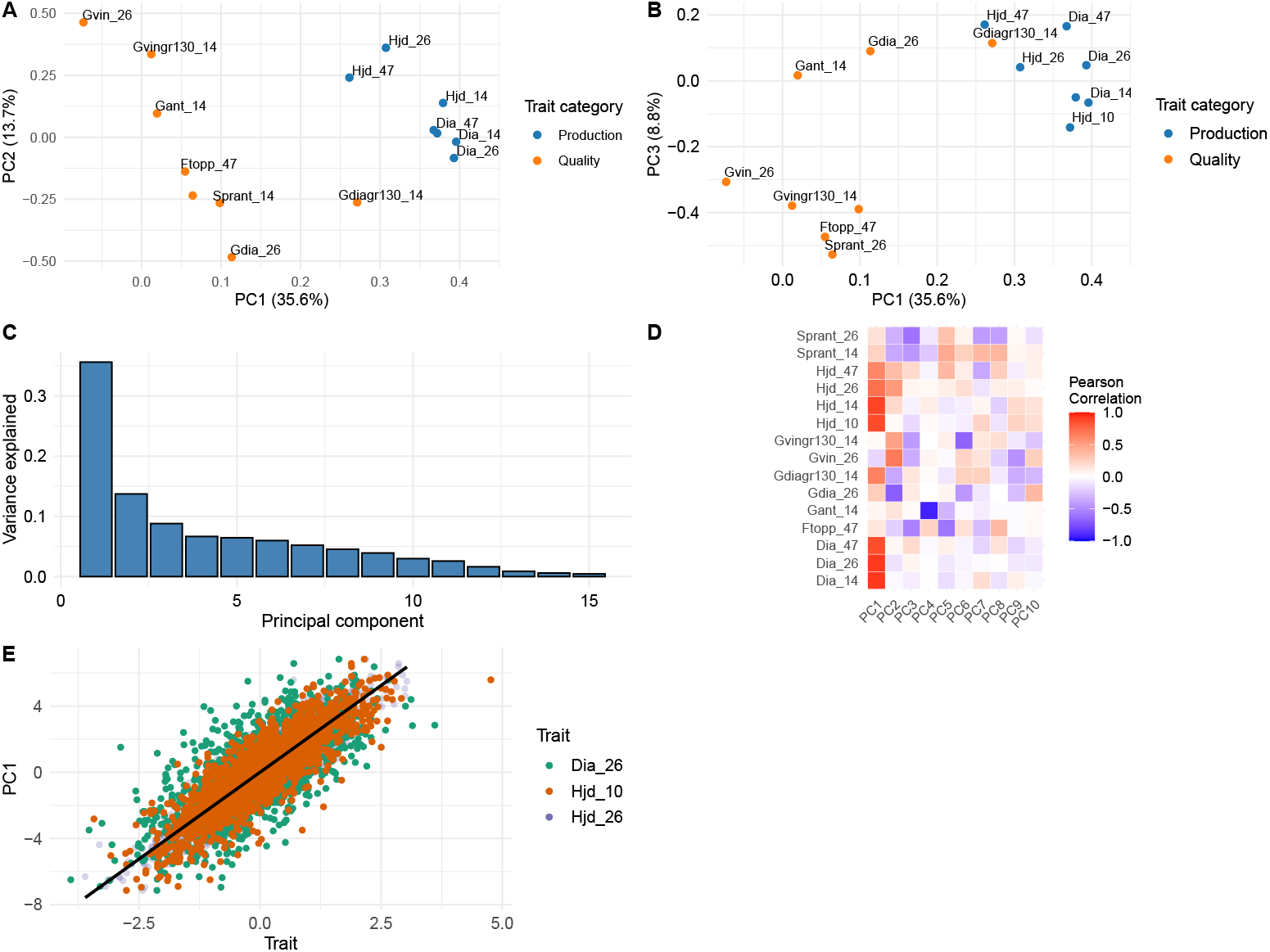
Principal component analysis on the 1685 tree subset with 15 traits measured: **A**, The loadings of PC one and two with trait categories colored, **B** loadings of PC one and three, **C** proportion of phenotypic variation explained by each of the 15 first PCs, **D** estimated Pearson correlation coefficient between original traits and the first ten PCs, and **E** scatter plot of PC1 scores and highest correlated trait values (Dia_26, Hjd_10, and Hjd_26).

The REML analysis on the original traits was carried out in several steps to make the model converge: a) a univariate REML analysis to estimate good starting values of the variance components, b) pairwise bivariate REML analysis of all trait combinations, and c) a BLUP analysis with fixed scale parameters obtained in step b). The estimated correlations and narrow-sense heritability 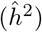 for all traits is shown in Fig 2 for both REML analyses of original and transformed traits. The range for 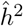 the original traits varied from 0.04 (Multiple stems year 47) to 0.57 (Height year 26) (Table 1). The computational time required for these steps where a) 2.51 (0.13) seconds, b) 94.3 seconds (individual runs ranging from 0.17 - 2.57 seconds, mean 0.89 (0.61) seconds), c) the model did not converge after 10 000 iterations (i.e. the log-likelihood maximum was not reached), although we deemed the model found a near optimal solution as the difference in the log-likelihood did not change in its third decimal across 10 iterations: as each iteration took 2.5 - 3.9 seconds, the total required time in the c) step was 7 hours, and 7 minutes.

**Figure 2.**
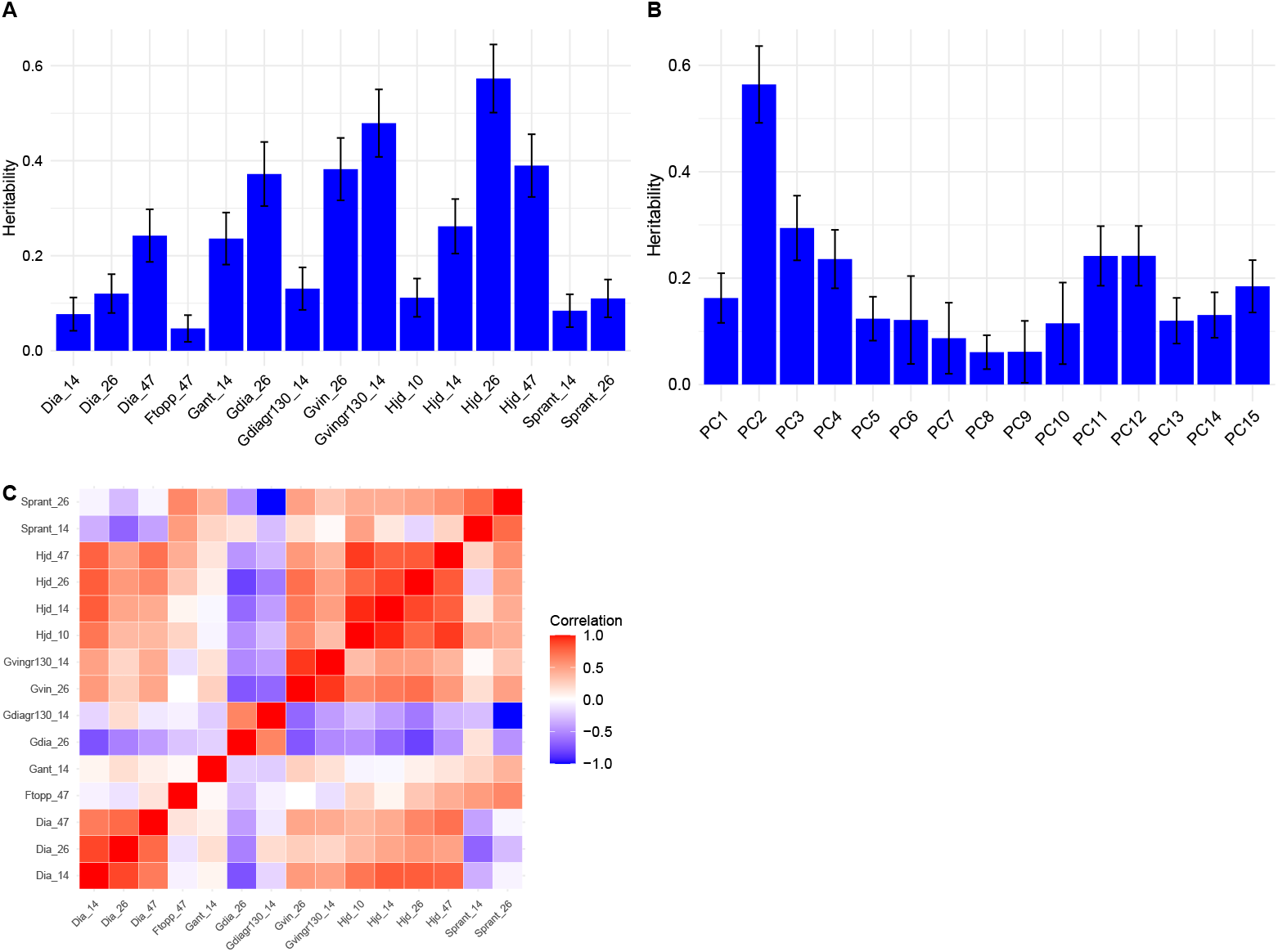
REML pairwise trait analysis on original traits and on PCA transformed traits (i.e. PCs) showing: **A** 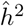 of the original 15 traits using both the bivariate and univariate approaches with standard errors displayed as vertical lines, **B** shows obtained 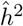 of the 15 PCs analysis, and **C** shows the pair-wise estimated correlations obtained with REML between all 15 traits.

Then, univariate REML analyses of all 15 PCs as response variables were performed as comparison, and in all cases converged after ten iterations in less than a second per PC. To verify that PCs are orthogonal and can be analyzed one by one, we analyzed PC1 and PC2 using bivariate REML:

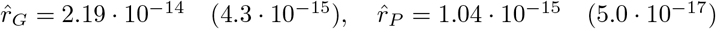, which imply independence between PC1 and PC2 both at genetic and phenotype levels. Obtained 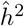 for all PCs are shown in Fig. 2B, and ranged between 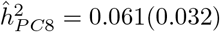 and 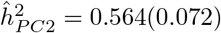, with standard error within parenthesis.

From a breeder’s perspective, the most crucial information is the ranking of EBVs and how one can utilize this information to perform selection and crossing (or mating) decisions. We defined two selection criteria: one production based with 50% EBVs for height and diameter at breast height at age 26, and one outer tree quality based with 50% production, 25% branch angle and negative 25% branch diameter, all measured at age 26. As a common breeding objective goal of forest trees is to increase the productivity, we included both height and diameter in both indices. In addition, outer tree quality traits, such as branch angle and branch diameter, will impact wood quality and their improvement are also important long term breeding goals. To make index based on PC traits comparable, we used Lasso shrinkage based regression to assign PCs to an index (Fig 3A), where most PCs does not contribute and three and four PCs contributed to production and quality index, respectively. In all, the correlation of obtained rank of the top 50 individuals based on selection index values from standard and PCA analysis were positive (Fig 3B and 3C: Kendall’s *τ* = 0.28, *p* = 0.0033, and *τ* = 0.44, 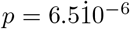, for production and quality index, respectively). Thus, although not identical, the rank lists resembled each other reasonably well in the top 50 rank, and similar response to selection is expected.

**Figure 3.**
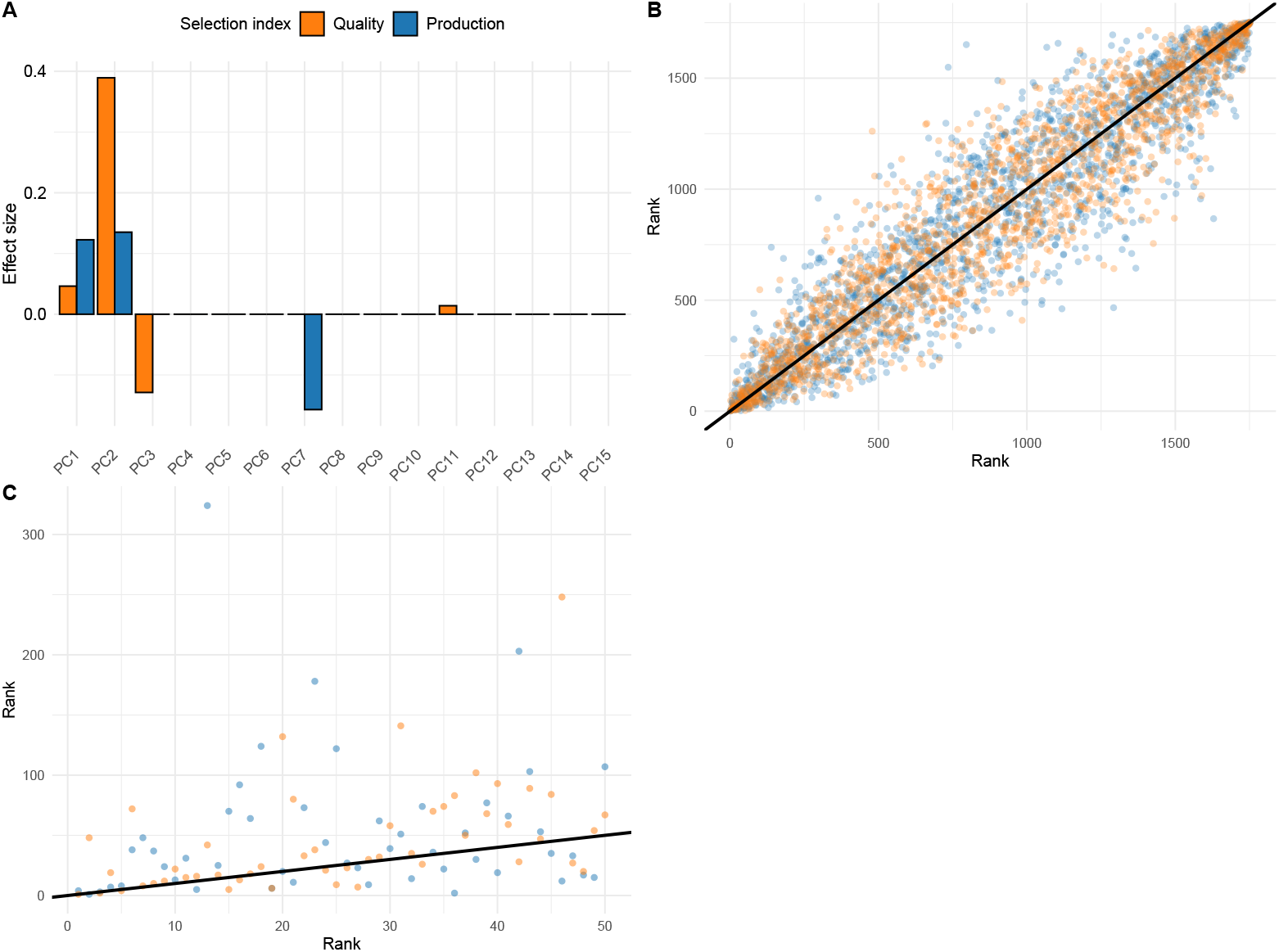
Comparison of tree rank based on estimated breeding values (EBVs) for PCA and standard approaches: **A** The contribution of individual PCs to the selection index corresponding to a traditional model both for quality and production based indices, **B** difference in rank of individuals where the rank of EBV based on analysis of original trait is the reference (x-axis) and the corresponding rank based on analysis of PC score traits (y-axis) for two selection index, **C** the same rank differences, but zoomed in on the first 50 reference ranked individuals. The color corresponds to the two selection indices in all sub-figures.

### Missing data can be efficiently handled

Standard SVD based PCA (SVD-PCA) cannot handle missing data. As data collected from realistic field trials would typically contain at least some proportion of missing observations, in particular if many traits have been measured. To check the effect of missing data on PCA based multi-trait selection, we selected a subset of 6044 trees scored for ten traits. The data contained 13.3% missing data in total ranging from 17 to 2021 missing observations for tree height at age 10 and age 47, respectively. The Bayesian PCA method (BPCA) was used to impute missing observations with nine PCs explaining 98.3% of the total variation.

To rule out that the imputed data had any impact on the genetic parameter estimates obtained by the REML method, we analyzed the data from 6044 individuals with 10 traits using the pairwise bivariate REML approach described earlier. We focus on the trait with the largest proportion of missing data (33.4%), tree height at age 47, as the worst-case scenario. The average difference in correlations to all other nine traits was 0.040 (0.035), with standard deviation within brackets. Correlations with some of the traits were overestimated with the imputed data, such as multiple stems and diameter at age 47, with 0.094 and 0.081, respectively. Obtained narrow-sense heritability estimates for the trait were 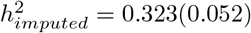 and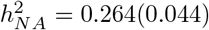.

However, this difference had little impact on EBV and ranking of trees as *r*_*Imp*,*NA*_ = 1.0, *p* = 2.2 × 10^*−*16^, probably because information is shared across correlated traits with much less fraction of missing data.

Furthermore, we investigated whether SVD based PCA (SVD-PCA) and BPCA yielded the same results for the same data. This was done in two steps: first, we checked the resulting ordinations to identify differences in the loadings and scores. Second, we estimated breeding values for the PCs generated with both SVD-PCA and BPCA to see if the rank of individuals differed. Both subsets were used: the 1685 tree subset with 15 traits without missing data, and the 6044 tree subset with 10 traits and imputed data. Distance matrices of the obtained loadings were highly correlated for both 1685 data (Mantel statistic: *r*_1685_ = 0.839, *p* = 0.001) and 6044 data (*r*_6044_ = 0.431, *p* = 0.006) (Supplementary figure S3). To investigate the impact on EBV (i.e. the scores of the ordinations), we calculated the EBV of PC1-PC3 with both methods on both data sets and calculated the Pearson correlations. All REML runs took 10 iterations to converge. EBV obtained with BPCA and SVD-PCA were highly correlated for all first three principal components, with all correlations obtained in the 1685 individual subset where equal to one, and slightly less than one for the 6044 individuals (*r*_*PC*1,6044_ = 0.997 (95% : 0.997, 0.998), *r*_*PC*2,6044_ = 0.998 (0.997, 0.999), and *r*_*PC*3,6044_ = 0.967 (0.966, 0.969)), with all *p <* 2.2 × 10^*−*16^ (Supplementary figure S4).

Taken together, there were barely noticeable differences between the BPCA and SVD-PCA approaches on the Scots pine progeny trial data, with the least difference at the EBV level of the PCs. In practice, both methods could be used interchangeably without changing the rank of trees. The need for imputing missing observations might be a bigger concern when it comes to estimating scale parameters but seems to be less important when considering rank of individuals based on EBV.

#### PCA allows for rapid multi-trait analysis in a *Pinus taeda* pedigree

In total, 27 traits were recorded for 926 individual genotypes in the South eastern USA breeding program of Loblolly pine (Table 2). 4. As missing data was present, we first removed traits with *>* 40% missing data, which resulted in the removal of the trait LesionUF 1 (i.e. damage to the tree that is caused by an unidentified factor after one growth season). In addition, individuals with *>* 25% missing data were removed, resulting in 861 pedigree member available for the multi-trait analysis. Missing data was imputed similarly as the Scots pine data example, with BPCA analysis of 20 PCs was performed explaining 97.1% of the total phenotypic variation. The BPCA analysis of the 26 traits revealed clustering tendencies (Fig 4A and B), with the first PC explaining 31.8% of the phenotypic variation, while the second and third explained 13.8% and 7.8% respectively. The first PC separates the production traits from the tree quality and disease susceptibility traits, albeit with some of the traits mixed (i.e. at PC1 values close to zero), such as branch diameter year 6 (BD 6), and the total tree height to the base of the live crown (HTLC 4). The second PC clearly separates production and tree quality traits from the disease susceptibility traits. The third PC separated crown width traits and branch diameter with various tree height traits.

**Table 2.**
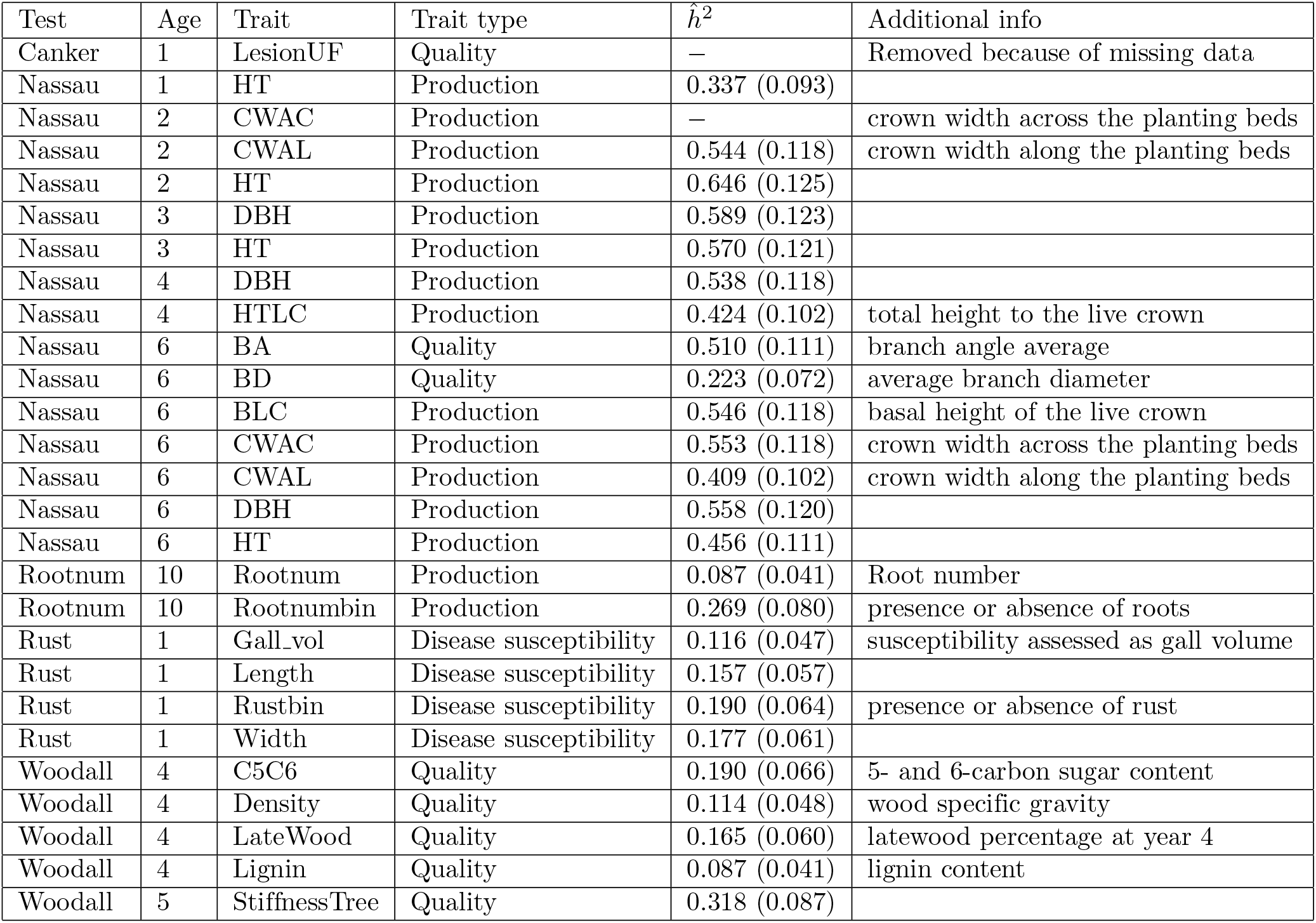
Loblolly pine trait statistics. Traits measured in the Lololly pine breeding population of 861 genotypes. Standard errors are within parenthesis. Estimates of narrow-sense heritability for each trait is denoted 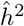.

**Figure 4.**
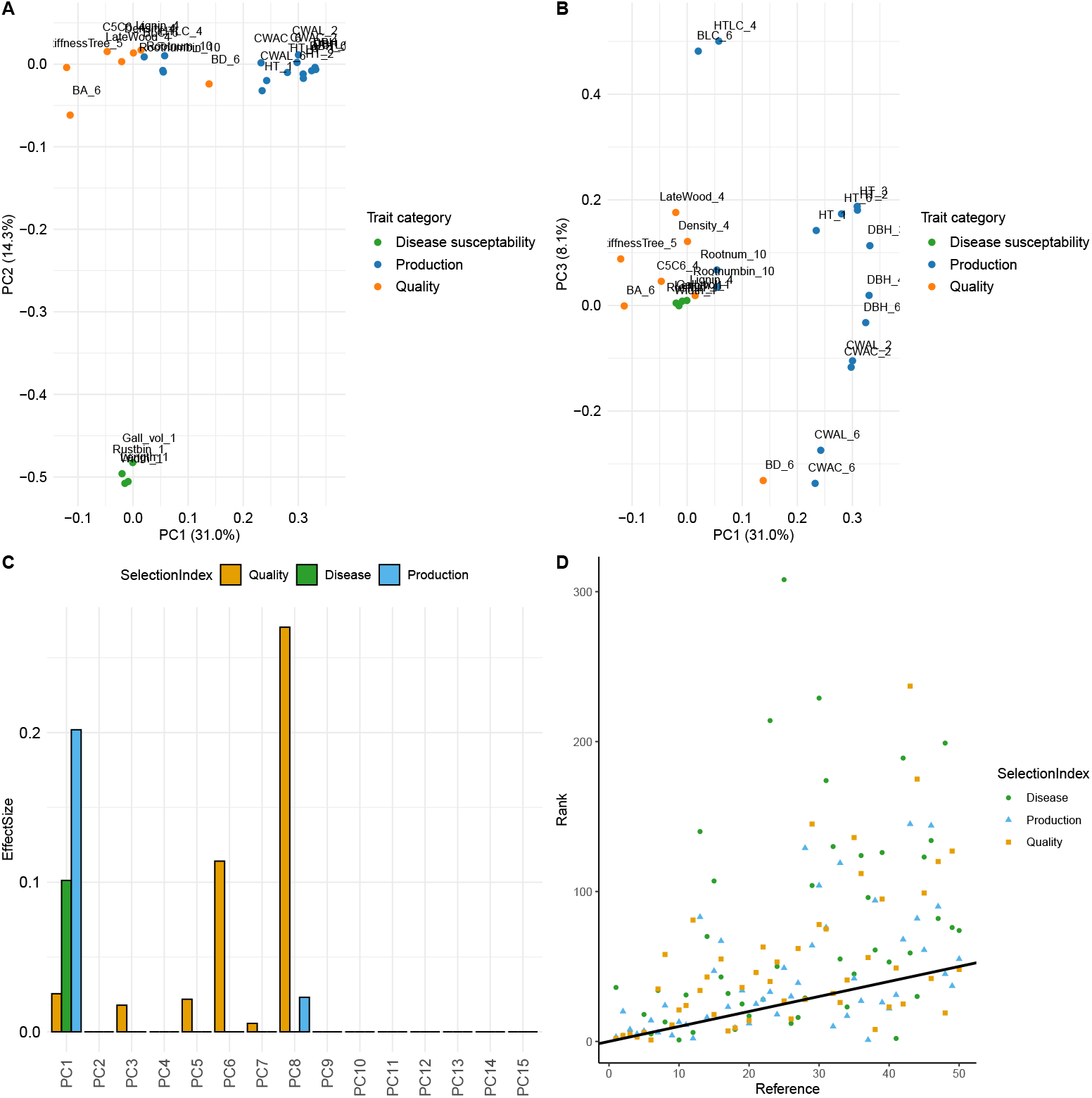
Multi-trait LMM analysis on the 26 selected traits in the Loblolly pine dataset. **A** Clustering of traits via PCs one and two colored with their respective trait category. The number on the axis labels corresponds to the percentage of phenotypic variation explained. **B** PC one and three. **C** Selected PCs for the three different selection indices and their respective effect size. **D** difference in rank of individuals where the rank of estimated breeding values (EBV) based on analysis of original trait is the reference (x-axis) and the corresponding rank based on analysis of PC score traits (y-axis) for three selection indices, zoomed in on the first 50 reference ranked individuals.

To test the impact of missing data imputation method, we also used the missForest method Stekhoven2012 and simple trait means to complete the data set and performed standard SVD-PCA (Supplementary figure S5). The BPCA explained 97.1% using 20 PCs with the first PC explaining 31.8%, while the missForest explained 98.7% in the first 20 PCs while the first PC explained 31.0%. When comparing the scores for the first 20 PCs in both BPCA and missForest imputed data resulted in highly correlated ordinations (Mantel’s *r* = 0.545, *p* = 0.001). Using trait means as imputation resulted in a very similar ordination as the missForest imputation SVD-PCA (results not shown). Thus, the method of imputation had relatively small impact on the resulting ordinations, although using the BPCA seemed to be advantageous if the first PC is of main interest (i.e. production traits).

To examine downstream results (i.e. rank lists) of PCA and pair-wise bivariate LMM approaches, we followed the same procedure as with the Scots pine example. First, to obtain the starting values of the bivariate REML analyses for estimating variance components, 26 univariate REML analyses were performed. In most cases, the model converged after 4-7 iterations, but in some cases, however, resulted in Log-likelihood not converging, and that some components changed by more than 1% on the last iteration. Each run was very quick, less than a second for all 26 traits. In total, 325 bivariate REML analyses were required to cover all trait combinations to estimate the trait covariance matrix. These analyses took 62 seconds in total, with very mixed convergence statistics ranging from four to 704 iterations. Finally, the BLUP analysis lasted for 12h 22min to run 3000 iterations until convergence, where each iteration took between 11 to 18 seconds.

To compare rank lists, we created three selection indices: one production-based index with 50% height and 50% diameter EBVs, one tree quality index combining height and diameter with tree density and wood stiffness all weighted equally, and finally a disease susceptibility index with height, diameter and fusiform rust presence. We used LASSO regularization to combine the PCs which best mimics these indices (Fig. 4C). The rank lists of the top 50 trees obtained with PCA and traditional approach resembled each other for all three selection indices considered (Fig. 4D): production index, Kendalls *τ* = 0.458, 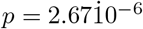, quality index Kendalls *τ* = 0.445, 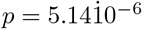, and disease susceptibility index, Kendalls *τ* = 0.468,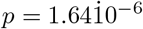. These results suggest that similar rank lists can be obtained with the PCA approach for three different indices but at a fraction of the required computing time.

To visualize the portability and flexibility of the approach, we tested a variety of available software implementations (Table 3). Thus, depending on the situation and requirement of the analysis and data, an analyst can choose among a large smorgasbord of alternatives.

**Table 3.**
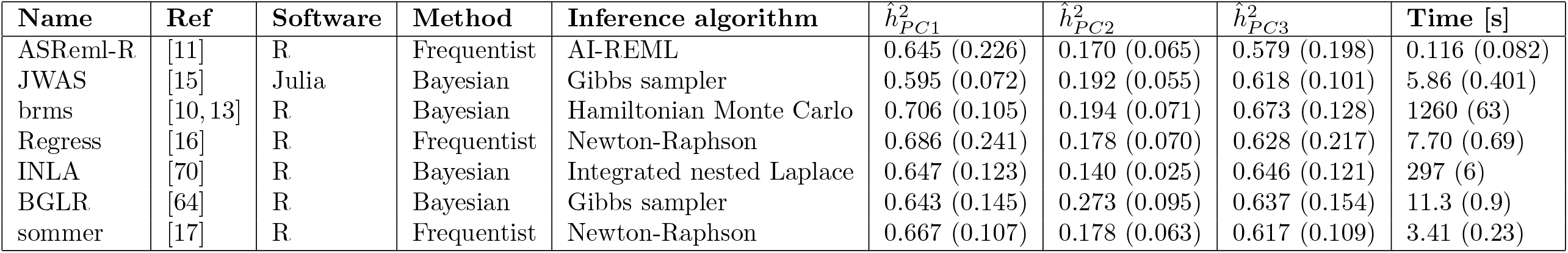
Linear mixed-effect model performance. Available linear mixed-effect model (LMM) implementations and their performance on the first three principal components in the Loblolly pine case study. Computing time is averaged across ten runs with standard deviations within parenthesis. All estimated heritabilities for each principal component (PC) are in the narrow-sense. Default settings was used.

| Name | Ref | Software | Method | Inference algorithm | $\hat{h}_{PC1}^2$ | $\hat{h}_{PC2}^2$ | $\hat{h}_{PC3}^2$ | Time [s] |
| --- | --- | --- | --- | --- | --- | --- | --- | --- |
| ASReml-R | [11] | R | Frequentist | AI-REML | 0.645 (0.226) | 0.170 (0.065) | 0.579 (0.198) | 0.116 (0.082) |
| JWAS | [15] | Julia | Bayesian | Gibbs sampler | 0.595 (0.072) | 0.192 (0.055) | 0.618 (0.101) | 5.86 (0.401) |
| brms | [10, 13] | R | Bayesian | Hamiltonian Monte Carlo | 0.706 (0.105) | 0.194 (0.071) | 0.673 (0.128) | 1260 (63) |
| Regress | [16] | R | Frequentist | Newton-Raphson | 0.686 (0.241) | 0.178 (0.070) | 0.628 (0.217) | 7.70 (0.69) |
| INLA | [70] | R | Bayesian | Integrated nested Laplace | 0.647 (0.123) | 0.140 (0.025) | 0.646 (0.121) | 297 (6) |
| BGLR | [64] | R | Bayesian | Gibbs sampler | 0.643 (0.145) | 0.273 (0.095) | 0.637 (0.154) | 11.3 (0.9) |
| sommer | [17] | R | Frequentist | Newton-Raphson | 0.667 (0.107) | 0.178 (0.063) | 0.617 (0.109) | 3.41 (0.23) |

## Discussion

Multi-trait LMM analysis to estimate heritabilities, genetic correlations and breeding values is the cornerstone of breeding programs for improving yield, disease resistance and quality in animals, crops and forest trees. Unfortunately, if many traits are considered jointly, this analysis is far from straight forward to perform due to several reasons, including problems with convergence to a stable solution, required computational time, and precision in parameter estimates. To overcome this hurdle, we propose the use of principal component analysis (PCA) to reduce the dimension of the phenotypic response variables. We show the benefit of the approach on two data set of Loblolly and Scots pine pedigrees, with a large number of traits recorded at multiple time points. The PCA separated trait groups and REML analysis resulted in an 1000 fold reduced computational time as compared to traditional multi-trait analysis. Because obtained principal components (PC) are othogonal (to each other), there is no need to use multivariate analysis and trait correlations are accounted for. The individual univariate REML analyses converged after 10 iterations. Rank lists based on estimated breeding values (EBV) obtained from traditional and PCA approaches correlated strongly among the different selection indices used (i.e. production, quality and disease resistance).

In breeding applications, it is not uncommon that the breeding objective traits cannot be measured directly, for example land economic value per hectare at the age of harvest in forest tree breeding programs [8]. In such cases, several traits are measured that hopefully correlate well with the breeding objective traits. This is a typical situation in breeding of species with long generation turnover, such as forest trees or some livestock animals. In these situations it could be more profitable to rather consider phenotypic profiles than individual traits with unclear connection to future breeding objective traits as diminishing age-age correlations reduces the response to selection [19, 41, 48]. As some traits are very expensive to measure, such as destructive sampling like meat quality traits in beef cattle [79], physiological traits in woody plants including fire-induced irreversible xylem damage and low temperature-induced tissue freezing [49], and wood (sawn timber) quality traits [26, 28], phenotype profiles could be measured and analysed with PCA to incorporate different types of traits jointly.

In Swedish forest tree breeding programs, there are currently multiple traits in assessment schemes including measurements on tree growth, adaptation and external wood quality. Similar characteristics have been incorporated into other tree breeding programs such as the fourth round of selection in the Loblolly pine breeding program in southeastern USA [39] and Douglas-fir breeding program in New Zeeland [21]. It is, however, expected that in the future the number of traits in selection will increase to further mitigate the effects of climate change on forest tree resilience and to aim for more adapted trees. Adaptation traits can be such as resistance to diseases and different pests [7, 33], spring frost tolerance [52], drought tolerance [35] and fecundity [49]. Furthermore, considering internal wood quality in terms of wood density measurement as a selection trait is under research development and is expected to have greater impact on breeding objectives in the future. Several studies have shown unfavourable genetic correlation between tree growth and wood density which should in that case take into account in breeding to maintain acceptable level of this trait for production purposes [27, 28]. Hence, the use of PCA based trait evaluations could drastically improve efficiency of multiple trait analysis, as both the PCA itself and the following univariate analyses can be conducted with great reduction in computing time without the loss of phenotypic and genetic variation.

In large-scale breeding evaluation systems, such as those provided by Interbull in dairy cattle (https://interbull.org/index), Treeplan in forest trees (http://www.treebreeding.com/technology/treeplan) and INGER in Rice (https://www.irri.org/inger), phenotyping and genotyping efforts are gathered and standardized on a wide geographical scale to perform selections for future generations of breeding. In such large scale programs, the genetic evaluation system play a crucial role in assessing the genetic merit of individuals. Data collected in trials with crops or forest trees typically need to be standardized, where site specific effects are removed from the phenotypic records, and genetic parameters must be collected at a population level to enhance nationwide or global comparison between available material. Then, depending on the breeding goal and target zone, all available trait data needs to be weighted together in the final BLUP analysis step. Thus, a number of analysis steps are conducted sequentially, to be able to evaluate all traits accordingly with reasonable accuracy and computing time. However, combining results from multiple PCA of different datasets is not straightforward because PCA is sensitive to the variance structure of the data it is applied to, and each analysis will reflect the unique variance structure (of that particular dataset). To circumvent this hurdle, harmonization of the data sets can improve such as pre-adjustment techniques to remove within site variation and set up common trait classes which should be included. In addition, incremental PCA (IPCA) [80] can be used as a feasible option to merge harmonized datasets into one very large. An alternative is to perform a meta-analysis of the results of each individual PCA to score which traits that are important for respective PC and identify common trends and ranklist similarities. [46] developed a sparse PCA alternative (MetaPCA) by combining the *L*1-regularization approach of [87] with a penalized matrix decomposition calculation, and showed improved accuracy in analysis of multiple omics datasets in yeast, prostate cancer, mouse metabolism and TCGA pan-cancer methylation. Further effort into this direction is needed to improve large scale genetic evaluations using PCA based methods.

Here, we used standard SVD based PCA and BPCA to obtain orthogonal PCs of all phenotypic traits. There are many alternative directions to improve this dimension reduction step, depending on the characteristics of the phenotypic data and the goal of the genetic evaluations. For example, each obtained PC in these example cases were a mixture of all included traits, albeit some to a very low degree. It is tempting simply to truncate small contributions of some variables, but Cadima1995 show that this ad-hoc solution can indeed result in erroneous approximations and poor interpretations. [87] introduced sparse PCA (SPCA) by performing a *L*1-regularization step via elastic nets so that sparse loadings is obtained, which greatly increases interpretability of the analysis (but see also [29, 45]). In a similar effort, [18] showed how to use simple or Hausman components to improve the interpretability of the analysis which satisfied the Thurstonian criteria (i.e., each component does not containing too many variables and each variable does not being incorporated into many components). These efforts could help in the genetic evaluations of breeding populations when creating selection indices for a more transparent use of PCs.

In the Scots pine example presented here, all traits were pre-adjusted prior to the ordination analysis to remove site specific effects [12], turning all the data as continuous traits, even though some were originally integer counts, such as the number of top shoots of the tree. Similarly, in the Loblolly pine example, all trait data (i.e. estimated breeding values) were adjusted or deregressed following the approach suggested by [30]. Continuous data works very well with PCA, as it relies on linear transformation that identifies the PC maximizing the variance in the data, regardless of the underlying distribution. However, the interpretation of the components is enhanced if the data is normally distributed. Non-normal data, especially if it includes outliers or is heavily skewed, can affect the estimation of the correlation matrix, which standard PCA relies on [44]. An alternative is robust PCA methods that are designed to handle data with outliers or noise that traditional PCA might not handle well: the method decomposes the data into a low-rank matrix and a sparse matrix, which can capture corrupted observations [29, 83]. In addition, there exist alternatives for non-continuous data, such as the multiple correspondence analysis (MCA), which is used for analyzing multivariate data sets containing categorical variables by creating an indicator matrix (a Burt table) from the original variables [58]. To summarize, there exists a smorgasbord of alternative PCA related approaches which can be used in situations of non-normal non-continuous data and to improve interpretability of PCA.

While PCA is best suited for continuous data, it is sometimes applied to discrete data in genetic analysis due to its popularity and ease of use. Widely used examples are applying PCA on binary marker data, such as SNPs or insertion-deletion (Indel) markers, to infer ancestral population assignments of the analyzed population or to correct for population stratification in genome wide association studies (GWAS), even though the discrete nature of the marker data violates the assumptions of the PCA. Some alternatives for overcoming this hurdle involves using correspondance analysis (i.e. MCA) or applying model based alternatives which can handle discrete data in analysis of genetic variation [2, 6]: further research into this direction is warranted.

In summary, we have shown that PCA can be a viable option in multi-trait analysis, and in particular if the number of traits measured is large. By reducing the multi-trait LMM to univariate alternatives, computing times can be 1000-fold reduced while capturing all phenotypic variation of the analyzed population. Several alternatives exists for data imputation to complete multi-trait records which allows for the use of PCA of phenotypic profiles in real breeding applications as highlighted in both of our case studies. Another advantage of the proposed approach is that there exists many available implementations of PCA and LMM which can be combined according to the specific application at hand. In our case, we tested some of the available LMM implementations (Table 3) that can accommodate various types of response functions, type of predictor sets and dependencies among those. We believe that PCA based genetic evaluations can be a part of a population genetic analysts toolbox for accurate and fast multi-trait analysis where large scale phenotyping efforts have been performed.

## Supporting Information

Project information including the Scots pine data and supplementary figures are provided at https://github.com/jonhar97/Reduced_phenotype_MME.

**S1 Figure**

**Observations per trait**. Number of observed phenotypic records for all spatially adjusted traits scored in the Scots pine field trial.

**S2 Figure**

**Trait dendrogram**. Dendrogram of the Euclidean distance between obtained loadings for: **A** the 1685 individual Scots pine subset with 15 traits available, and **B** the 6044 individual subset with 10 traits available. Blue branches highlights production related traits while orange branches highlights tree quality related traits.

**S3 Figure**

**BPCA - SVD-PCA comparison**. Loadings of principal component one (PC1) and PC2 obtained for Bayesian PCA (BPCA) and SVD-PCA for: **A** complete 1685 observations without missing data, and **B** for 6044 observations with imputed data using BPCA.

**S4 Figure**

**EBV comparison**. Estimated breeding values (EBV) of principal components PC1, PC2 and PC3 obtained using BPCA and SVD-PCA for **A** − **C** the 1685 Scots pine data set and **D** − **F** the 6044 data set.

**S5 Figure**

**Missing data imputation comparison**. Loading plot of PC1 and PC2 on the Loblolly pine data using the three tested imputation methods BPCA, missForest and trait average.

## Acknowledgments

This work was supported by the Trees4Future project (https://www.slu.se/centrumbildningar-och-projekt/trees-and-crops-for-the-future/t4f/t4f_intro2/) for JA, DH and MS.

## Notes

https://github.com/jonhar97/ReducedphenotypeMME

### Competing Interest Statement

The authors have declared no competing interest.

