## Supplementary figures and images for "Principal component analysis revisited: fast multi-trait genetic evaluations with smooth convergence"

### Supplemental Figure 1

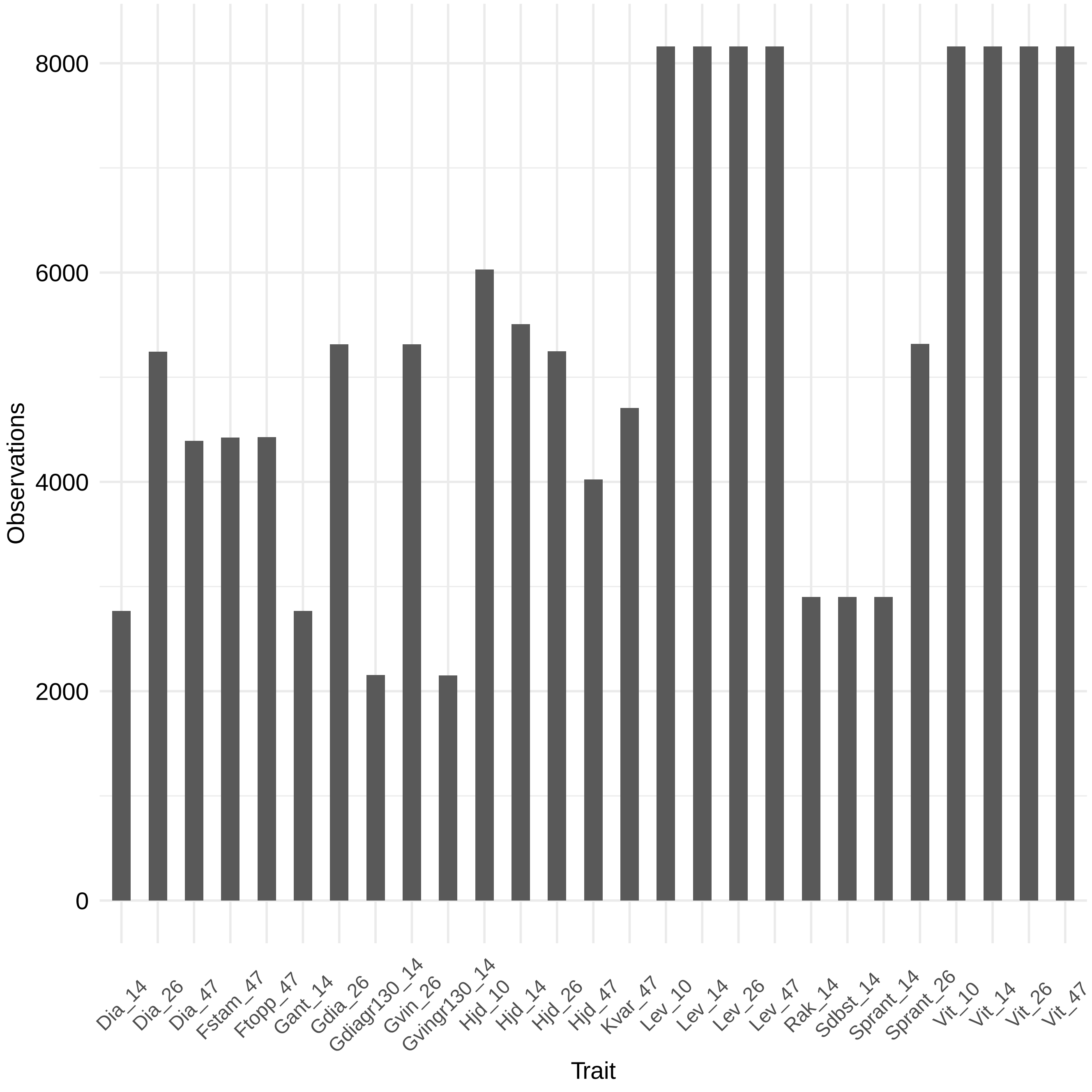

### Supplemental Figure 2

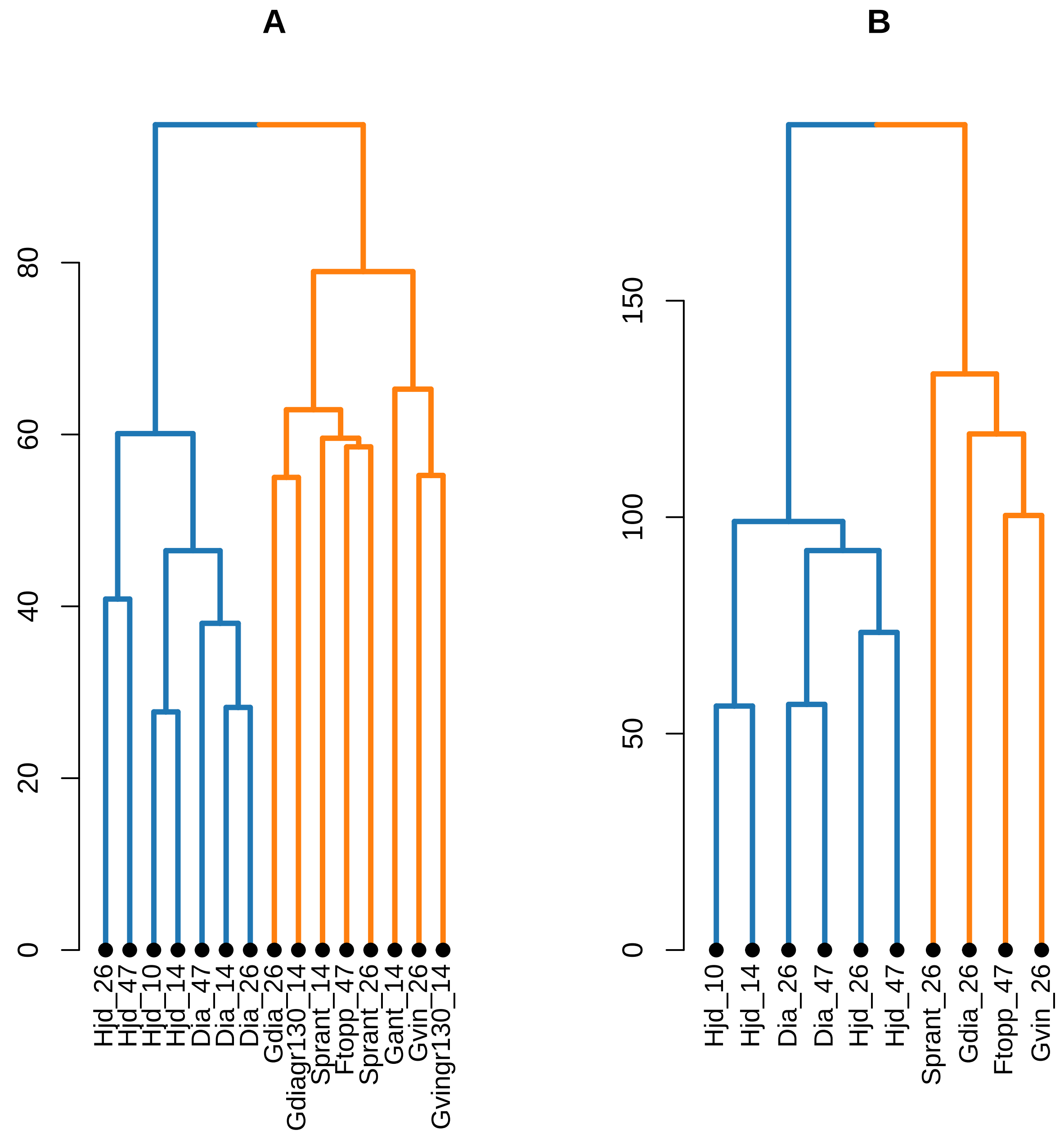

### Supplemental Figure 3

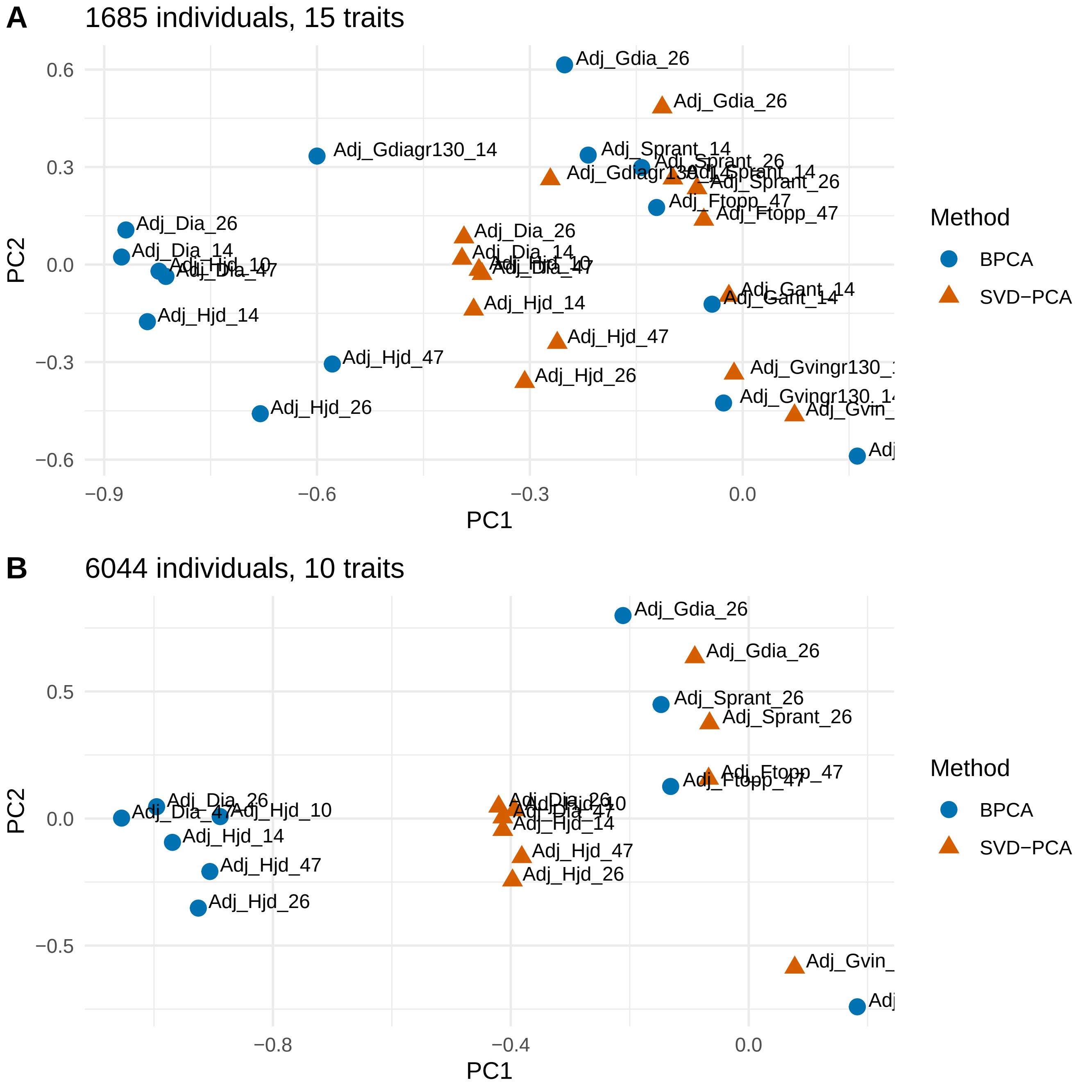

### Supplemental Figure 4

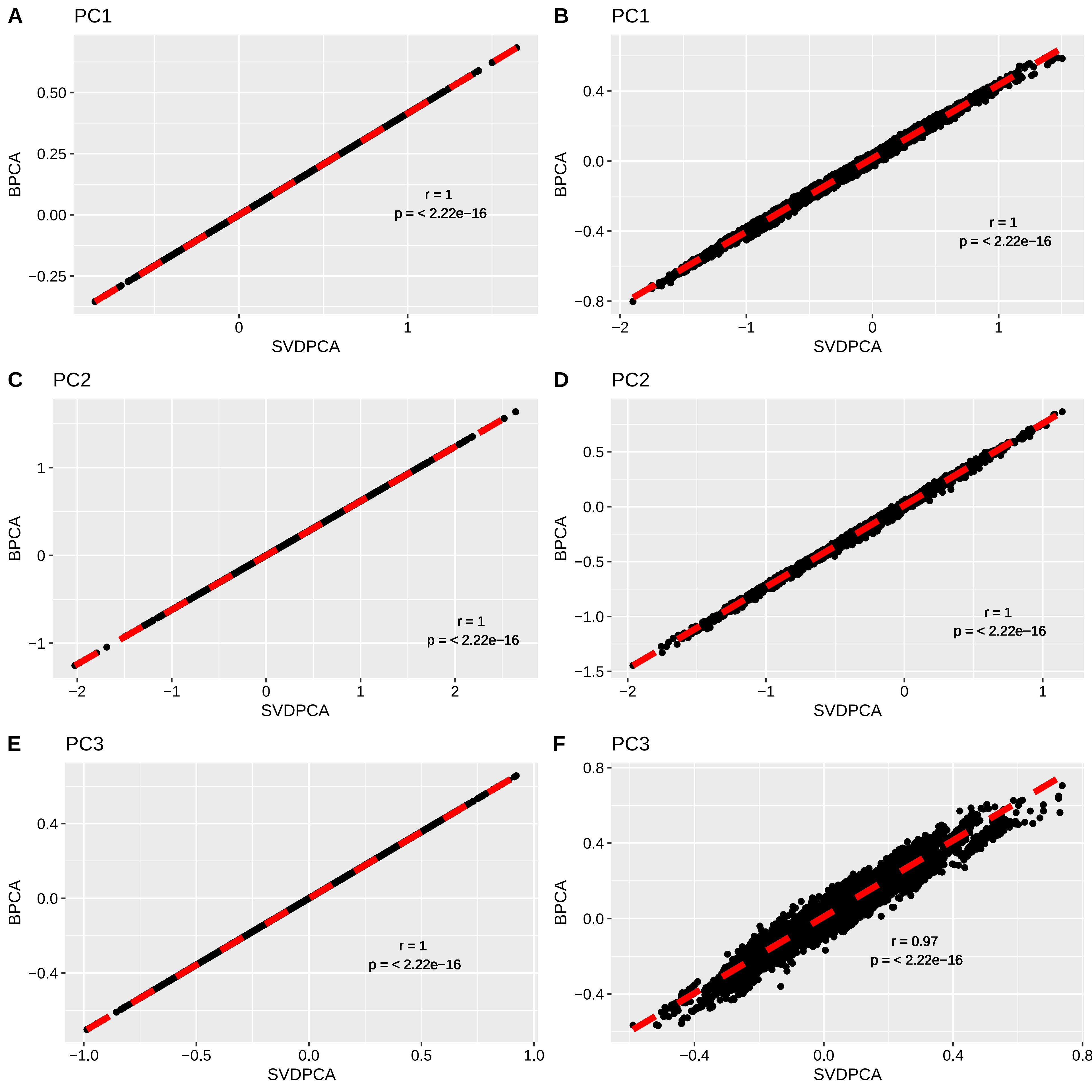

### Supplemental Figure 5

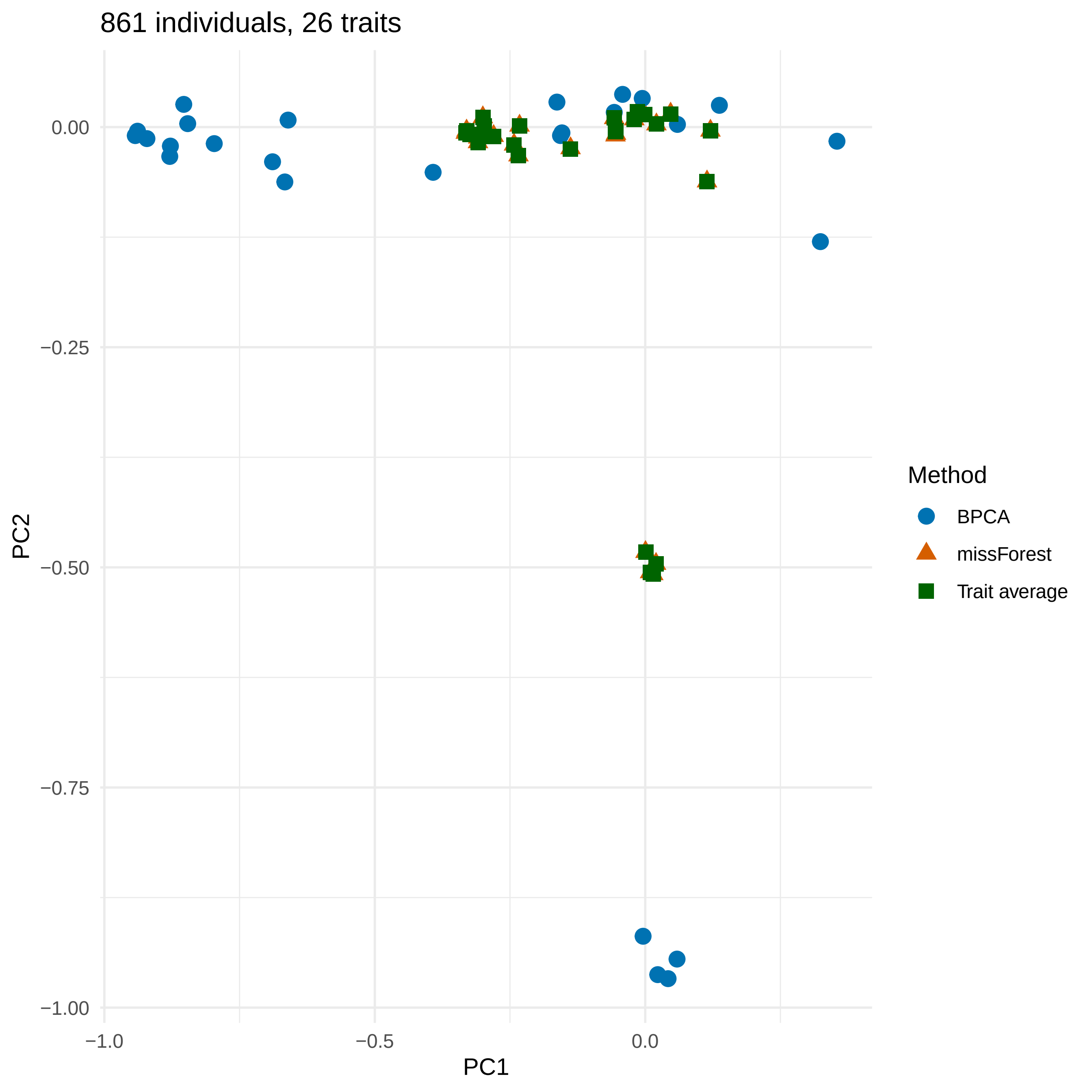
